# A reduction methodology for fluctuation driven population dynamics

**DOI:** 10.1101/2021.01.28.428565

**Authors:** Denis Goldobin, Matteo di Volo, Alessandro Torcini

## Abstract

Lorentzian distributions have been largely employed in statistical mechanics to obtain exact results for heterogeneous systems. Analytic continuation of these results is impossible even for slightly deformed Lorentzian distributions, due to the divergence of all the moments (cumulants). We have solved this problem by introducing a *pseudo-cumulants’* expansion. This allows us to develop a reduction methodology for heterogeneous spiking neural networks subject to extrinsinc and endogenous fluctuations, thus obtaining an unified mean-field formulation encompassing quenched and dynamical disorder sources.

## Introduction

The Lorentzian distribution (LD) is the second most important stable distribution for statistical physics (after the Gaussian one) [1], expressible in a simple analytic form, i.e.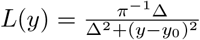, where *y*_0_ is the peak location and Δ is the half-width at half-maximum (HWHM). In particular, for a system with random heterogeneities distributed accordingly to a LD the average observables can be estimated exactly via the residue theorem [2].

This approach has found large application in physics, ranging from quantum optics, where it was firstly employed to account exactly for the presence of heterogeneities in laser emission [2, 3], to condensed matter, where by assuming a LD for the potential disorder, within the Lloyd model [4], it was possible to obtain exact results for the Anderson localization in three dimensions [5]. Furthermore, thanks to a Lorentzian formulation exact results can be obtained for the collective dynamics of heterogeneous oscillators in various contexts [6, 7]. Moreover, LDs emerge naturally for the phases of self-sustained oscillators driven by common noise [8, 9].

More recently, the Ott–Antonsen (OA) Ansatz [10, 11] yielded closed mean-field (MF) equations for the dynamics of the synchronization order parameter for globally coupled phase oscillators on the basis of a wrapped LD of their phases. The nature of these phase elements can vary from biological and chemical oscillators [12, 13] through superconducting Josephson junctions [14, 15] to directional elements like active rotators [16, 17] or magnetic moments [18].

A very important recent achievement has been the application of the OA Ansatz to heterogeneous globally coupled networks of spiking neurons, namely of quadratic integrate-and-fire (QIF) neurons [19, 20]. In particular, this formulation has allowed to derive a closed low-dimensional set of macroscopic equations describing exactly the evolution of the population firing rate and of the mean membrane potential [21]. In the very last years the Montbrió–Pazó–Roxin (MPR) model [21] is emerging as a representative of a new generation of neural mass models able to successfuly reproduce relevant features of neural dynamics [22–33].

However, the OA Ansatz (as the MPR model) is unable to capture the role of random fluctuations naturally present in real systems. In brain circuits the neurons are sparsely connected and *in vivo* the presence of noise is unavoidable [34]. These different sources of fluctuations are at the origin of fundamental aspects of neural dynamics, as the balance among excitation and inhibition [35, 36] and the emergence of collective behaviours [37]. The inclusion of these ingredients in MF models has been so far limited to homogeneous neural populations [38–43]. The main scope of this Letter is to fill such a gap by developing a unified MF formalism for heterogeneous noisy neural networks encompassing quenched and dynamical disorders.

In more details, in this Letter we introduce a general reduction methodology for dealing with deviations from the LD on the real line. This approach is based on the expansion of the characteristic function in terms of *pseudo-cumulants*, thus avoiding the divergences related to the expansion in conventional moments or cumulants. The implementation and benefits of this formulation are demonstrated for heterogenous populations of QIF neurons in presence of extrinsic and endogenous noise sources, where the conditions for a LD of the membrane potentials [21] are violated as in [44, 45]. In particular, we will derive a hierarchy of low-dimensional MF models for noisy globally coupled populations and deterministic sparse random networks, with a particular emphasis on the emergence of fluctuation driven collective oscillations (COs). For all these realistic setups we will show that our formulation reproduces quantitatively the network dynamics, while the MPR model fails in giving even a qualitative picture.

### Heterogeneous populations of QIF neurons

Let us consider a globally coupled network of *N* heterogeneous QIF neurons, whose membrane potentials {*V_j_*}, with *j* = 1,…, *N*, evolve as

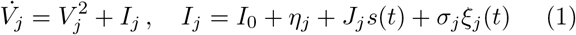

where *I*_0_ is the external DC current, *η_j_*: the neural excitability, *J_j_s(t)*: the recurrent input due to the neural activity *s(t)* mediated by the synaptic coupling *J_j_*. Furthermore, each neuron is subject to an additive Gaussian noise of amplitude *σ*_*j*_ = *σ*(*η_j_*, *J_j_*), where 〈ξ_*j*_(*t*)ξ*_l_*(*t*′)〉 = 2δ_*jl*_δ(*t* − *t*′) and 〈ξ_*j*_〉 = 0. The *j*-th neuron emits a spike whenever the membrane potential *V_j_* reaches +∞ and it is immediately reset at −∞ [46]. For instantaneous synapses, in the limit *N* →∞ the activity of the network *s(t)* will coincide with the population firing rate *r(t)* [21]. Furthermore, we assume that the parameters *η_j_* (*J_j_*) are distributed accordingly to a LD *g*(*η*) (*h(J)*) with median *η*_0_ (*J*_0_) and HWHM Δ_*η*_ (Δ_*J*_).

In the thermodynamic limit, the population dynamics can be characterized in terms of the probability density function (PDF) *w*(*V, t|**x***) with ***x*** = (*η*, *J*), which obeys the following Fokker–Planck equation (FPE):

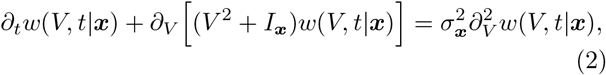

where ***I***_***x***_ ≡ *I*_0_ + *η* + *Jr(t)*. In [21], the authors made the Ansatz that for any initial PDF *w*(*V*, 0|***x***) the solution of Eq. (2) in absence of noise converges to a 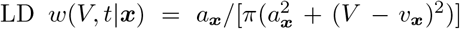, where *v*_***x***_ and 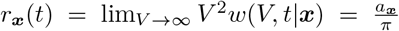 represent the mean membrane potential and the firing rate for the ***x***-subpopulation. This Lorentzian Ansatz has been shown to correspond to the OA Ansatz for phase oscillators [21] and joined with the assumption that the parameters *η* and *J* are indipendent and Lorentzian distributed lead to the derivation of exact low dimensional macroscopic evolution equations for globally coupled deterministic QIF networks.

### Characteristic function and pseudo-cumulants

In order to extend the MPR approach [21] to noisy systems, we introduce the characteristic function for *V*_***x***_, i.e. the Fourier transform of its PDF, namely

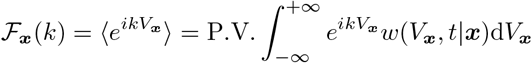

in this framework the FPE (2) can be rewritten as

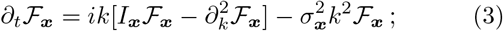

for more details see [47]. Under the assumption that 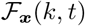 is an analytic function of the parameters ***x*** one can estimate the average chracteristic function for the population 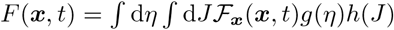 and the corresponding FPE via the residue theorem, with the caution that different contours have to be chosen for positive (upper half-plane) and negative *k* (lower half-plane).

Hence, the FPE is given by

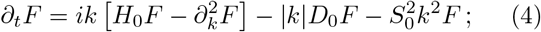

where *H*_0_ = *I*_0_ + *η*_0_ + *J*_0_*r*, *D*_0_ = Δ_*η*_ + Δ*_J_r* and 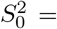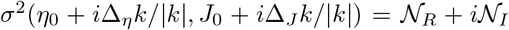. For the logarithm of the characteristic function, *F(k)* = Φ^*(k)*^, one obtains the following evolution equation :

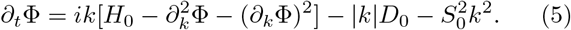

In this context the Lorentzian Ansatz amounts to set Φ_*L*_ = *ikv* − a|*k*| [48], by substituting Φ_*L*_ in (5) for *S*_0_ = 0 one gets

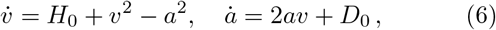

which coincides with the two dimensional MF model found in [21] with *r* = *a*/π.

In order to consider deviations from the LD, we analyse the following general polynomial form for Φ

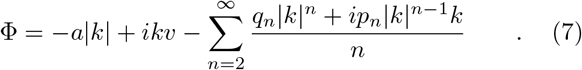

The terms entering in the above expression are dictated by the symmetry of the characteristic function 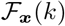 for real-valued *V*_***x***_, which is invariant for a change of sign of *k* joined to the complex conjugation. For this characteristic function neither moments, nor cumulats can be determined [49]. Therefore, we will introduce the notion of *pseudo-cumulants*, defined as follows

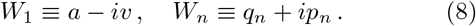

By inserting the expansion (7) in the Eq. (5) one gets the evolution equations for the pseudo-cumulants, namely:

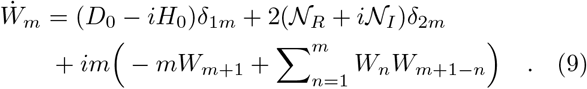

It can be shown [47] that the modulus of the pseudo-cumulants scales as |*W_m_*| ∝ |*S*_0_|^2(*m*−1)^ with the noise amplitude, therefore it is justified to consider an expansion limited to the first two pseudo-cumulants. In this case, one obtains the following MF equations

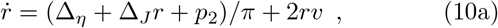

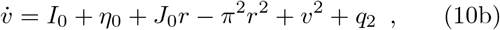

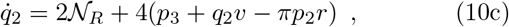

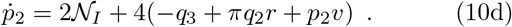

As we will show in the following this four dimensional model (with the simple closure *q*_3_ = *p*_3_ = 0) is able to accurately reproduce the macroscopic dynamics of noisy globally coupled and deterministic sparse QIF networks. Therefore the MF model (10) represents an extention of the MPR model to system subject to extrinsic and/or endogenous noise sources.

As shown in [47], the definitions of *r* = lim_*V*→∞_ *V*^2^*w(V, t)* and 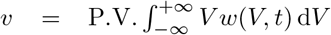 in terms of *w(V, t)* reported in [21] for the LD are not modified by considering corrective terms {*q_n_*, *p_n_*} of any order. Furthermore, *q*_2_ (*p*_2_) can be interpreted as an analougue of *kurtosis* (*skenwess*) for a distorted Gaussian distribution [47].

### Globally coupled network with extrinsic noise

To show the quality of the MF formulation (10) let us consider a globally coupled network of QIF neurons each subject to an independent additive Gaussian noise term of amplitude *σ* (i.e. 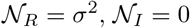. In this setup, the model (10) is able to reproduce the macroscopic dynamics of the network in different dynamical regimes relevant for neural systems.

Let us first consider the asynchronous dynamics, this corresponds to a fixed point solution 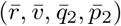 for (10). As shown in Fig. 1 (a-b), in the asynchronous state (AS) the MF model (10) reproduces quite well the population firing rate and the mean membrane potential obtained by the network simulations, while the deviations from the MPR model (dashed magenta lines) become appreciable for noise amplitudes 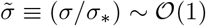, where *σ*_*_ represents a reference noise scale defined in [47]. Furthermore, the corrections *q*_2_ and *p*_2_ scales as ∝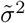 as expected (see Fig. 1 (c-d)). The truncation to the second order of the expansion (9), which leads to (10), is largely justified in the whole range of noise amplitude here considered. Indeed as displayed in Fig. 1 (e) 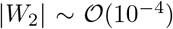 and 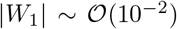, while the moduli of the other pseudo-cumulants are definitely smaller, more details in [47].

**FIG. 1.**
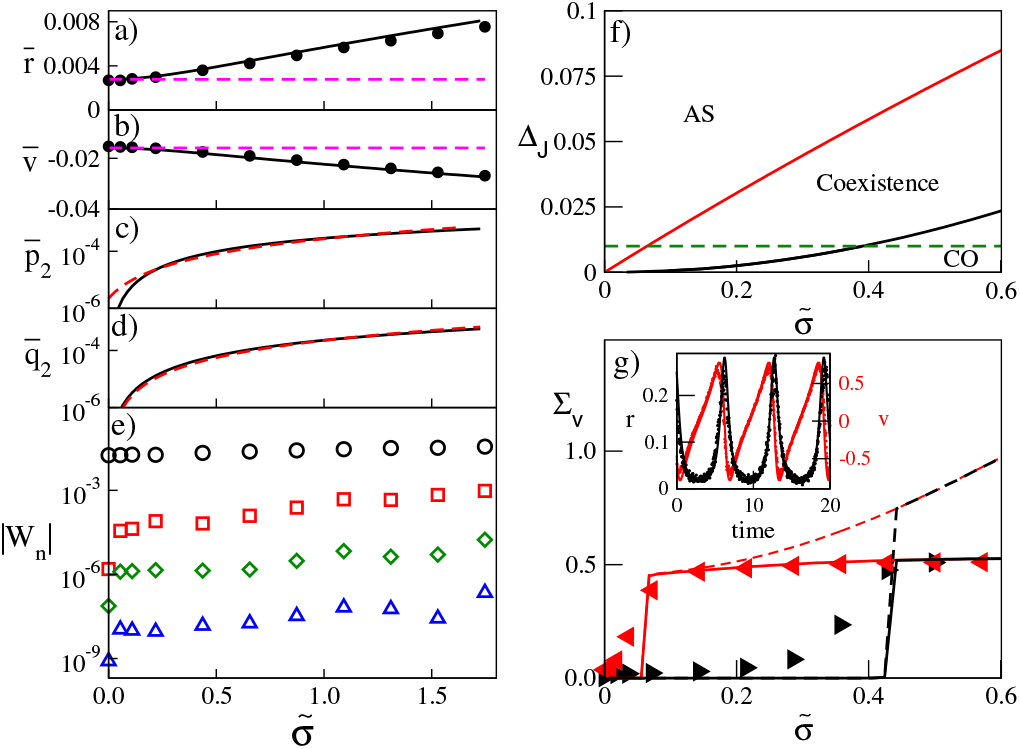
(a-e) Asynchronous Dynamics. Stationary values 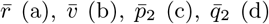 and |*W_n_*| (e) versus the rescaled noise amplitude 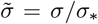 [47]. (a-b) Symbols refer to network simulations with *N* = 16000, solid line to the MF model (10), dashed (magenta) lines are the values of 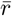 and 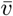 for the MPR model. In (c-d) the dashed red lines refer to a quadratic fit to the data. In (e) the symbols from top to bottom denote |*W*_1_|, |*W*_2_|, |*W*_3_| and |*W*_4_|. Other parameters : *I*_0_ = 0.0001, *J*_0_ = −0.1, Δ_*J*_ = 0.1, *σ*_*_ = 0.00458. **(f-g) Emergence of COs**(f) Bifurcation diagram in the plane 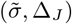 : the black (red) solid line denotes sub-critical Hopf bifurcations from AS to COs (saddle-node bifurcations of limit cycles). The horizontal green dashed line corresponds to the case studied in (g). (g) Standard deviations ∑_*v*_ obtained for quasi-adiabatic variation of 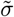. Lines (symbols) refer to MF (network) simulations: black (red) lines and right (left) triangles are obtained by increasing (decreasing) *σ*. Dashed (solid) lines refer to the pseudo-cumulant reduction (9) arrested to the second (third) order. In the inset are reported *r* and *v* versus time for *σ* = 0.143: dots refer to network simulations with *N* = 32000 and lines to MF results. Other parameters: *I*_0_ = 0.38, *J*_0_ = −6.3, Δ_*J*_ = 0.01, *σ*_*_ = 0.014, *η*_0_ = Δ_*η*_ = 0.

For lower levels of heterogeneity, one can observe the emergence of noise induced COs. The analysis of the bifurcation diagram of the MF model (10) in the plane 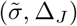 reveals the existence of three dynamical states: asynchronous, oscillatory and a regime of coexistence of AS and COs. In particular, a large heterogeneity Δ_*J*_ prevents the occurrence of COs, which are instead promoted by strong noise. As shown in Fig. 1 (f), the regimes are delimited by bifurcation lines: namely, COs emerge via sub-critical Hopf bifurcations from AS (black solid line) and disappear via a saddle-node bifurcation of limit cycles (red solid line). The dynamical regimes induced by noise cannot be captured by the MPR model, which would predict only the AS corresponding to 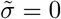.

Let us focus on a specific cut in the plane 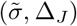, namely we fix Δ_*J*_ = 0.01 (green dashed line in Fig. 1 (f)). In this case the MF reveals that the AS looses stability in favour of noise driven COs at 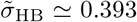 due to a sub-critical Hopf bifurcation, and that COs can survive down to 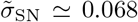, where they disappear via a saddle-node bifurcation. These transitions can be characterized in terms of the standard deviation ∑_*v*_ of the mean membrane potential, which is zero (finite) in the AS (CO regime) in the thermodynamic limit. We report in Fig. 1 (g) ∑_*v*_ versus 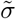 as obtained from adiabatic simulations of the MF (solid and dashed lines) and of the network with *N* = 64000 (triangles). Apart finite time and size effects that prevent the vanishing of ∑_*v*_ in the AS, the network simulations reveal the same dynamical behaviours as the MF. Furthermore, as shown in the inset of panel (*g*) the model (10) is able to accurately reproduce the time evolution of *v* and *r* also during COs for moderate noise amplitudes (namely, 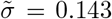. However, at larger noise amplitudes in order to obtain a good quantitative agreement between MF and network simulations one should extend the pseudo-cumulant reduction (9) to the third order (solid lines in panel (g)).

### Sparse networks exhibiting endogenous fluctuations

Let us now consider a sparse random network characterized by a LD of the in-degrees *k*_*j*_ with median *K* and HWHM Δ_*k*_ = Δ_0_*K*, a scaling assumed in analogy with exponential distributions. By following [38], we can assume at a MF level that each neuron *j* receives *k_j_* Poissonian spike trains characterized by a rate *r*, this amounts to have an average synaptic input 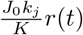 with Gaussian fluctuations of variance 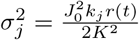. As shown in [44, 50], the quenched disorder in the synaptic inputs can be rephrased in terms of heterogeneous synaptic couplings. Namely, we can hypothesise for the MF formulation that the neurons are fully coupled, but with random distributed synaptic couplings 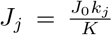 with median *J*_0_ and HWHM Δ_*J*_ = |*J*_0_| Δ_0_. Furthermore, each neuron *j* will be subject to a noise of intensity 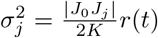 and this amounts to have 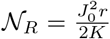 and 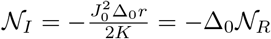.

For this random network, we report in Fig. 2 (a) a bifurcation diagram in the plane (|*J*_0_|, *I*_0_) estimated for the MF model (10). We find the same three dynamical regimes observed in the globally coupled case, but differently arranged and separated by different bifurcation lines. In particular, the AS emerges for sufficiently large excitatory drive *I*_0_, where the most part of neurons are supra-threshold and the dynamics is essentially mean driven [51]. The coupling strenght controls the amplitude of current fluctuations, indeed for increasing |*J*_0_| fluctuation driven COs emerge via a super-critical Hopf bifurcations (black solid line). For even larger coupling, beyond a sub-critical Hopf bifurcation (red solid line) a coexistence among AS and CO can be observed. However, the nature of this AS is different from that at low |*J*_0_|: this is a fluctuation driven regime, where excitatory drive and recurrent inhibition tend to balance [35, 36].

**FIG. 2.**
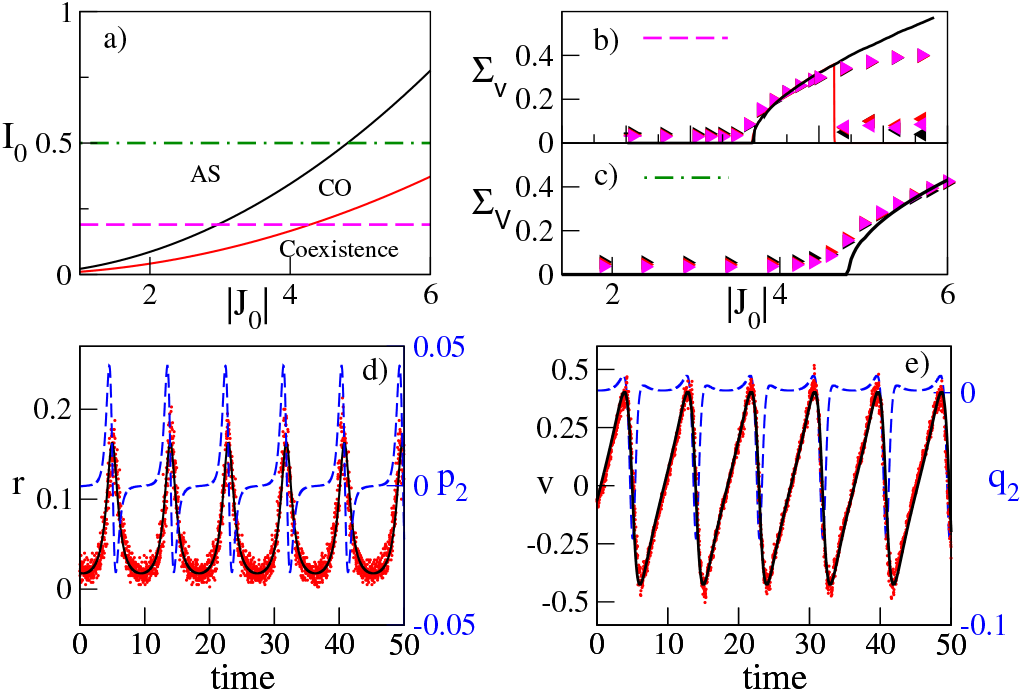
Sparse Networks. (a) Biurcation diagram in the plane (|*J*_0_|, *I*_0_): the black (red) line denotes super-critical (sub-critical) Hopf bifurcations from AS to COs. The magenta dashed (green dot-dashed) line indicates the case examined in panel (b) for *I*_0_ = 0.19 (panel (c) for *I*_0_ = 0.50). (b-c) ∑_*v*_ obtained for quasi-adiabatic variation of |*J_0_*|. Solid line (colored symbols) refers to MF (network) results: black (red) lines and right (left) triangles are obtained by increasing (decreasing) |*J*_0_|. The symbol colors denote different network sizes: namely, *N* = 10000 (black), 15000 (red), and 20000 (magenta). The values are averaged over 8 random realisations of the network. For *J*_0_ = 3.7 and *I*_0_ = 0.19 the MF evolution of *r* (*p*_2_) and *v* (*q*_2_) is displayed in (d) and (e) as black solid (blue dashed) lines, respectively. In (d-e) the symbols refer to network simulations with *N* = 40000. In all panels the parameters are *K* = 4000, Δ_0_ = 0.01, *η*_0_ = Δ*η* = 0.

As shown in Fig. 2 (b-c), network simulations are in good agreement with the MF predictions for low and high DC currents, as it can be appreciated by comparing the standard deviations ∑_*v*_ obtained for network of different sizes with the findings of the model (10). Furthermore, at intermediate coupling the oscillations of *r* and *v* in the network are very well captured by the MF dynamics (see Figs. 2 (d) and (e)). However, as shown in panel (b) for *I*_0_ = 0.19 the amplitudes of the COs are slightly overestimated by the MF at sufficiently large |*J*_0_| > 4.5. No discrepancies are observable in panel (c) for the same range of |*J*_0_|. As shown in [47] this is due to the fact that the rescaled noise amplitudes are definitely smaller in this latter case.

As shown in [44], the MPR model, even with the inclusion of the quenched disorder due to the heterogeneous in-degrees, is unable to predict the oscillatory and coexistence regimes displayed by the sparse inhibitory network. Therefore, it is fundamental to take in account corrections to the Lorentzian Ansazt due to endogenous fluctuations. Indeed, as shown in Fig. 2 (d) and (e) the evolution of *r* and *v* is clearly guided by that of the corrective terms *p*_2_ and *q*_2_ displaying regular oscillations.

## Conclusions

fundamental aspect that renders the LD difficult to employ in a perturbative approach is that all moments and cumulants diverge. However, to cure this aspect one can introduce an expansion in *pseudo-cumulants* of the characteristic function. As we have shown, this expansion can be fruitfully applied to build in full generality a hierarchy of low-dimensional neural mass models able to reproduce, with the desidered accuracy, firing rate and mean membrane potential evolutions for heterogeneous spiking neural networks in presence of extrinsic and intrinsic fluctuations.

One of the main important aspects of the MPR formulation, as of our reduction methodology, is the ability of these MF models to capture transient synchronization properties and oscillatory dynamics present in the spiking networks [22, 26, 27, 32], but that are lost in usual rate models as the Wilson-Cowan one [52]. However, our MF formulation can encompass further fundamental features of brain circuits beyond heterogeneity, as sparsness in the synaptic connections and background noise [34], not envisaged in the MPR model. Therefore, our mass model is able to reproduce spiking network dynamics induced by various noise sources such as irregular firing regimes or fluctuation driven COs, which cannot be predicted within the MPR framework developed for globally coupled deterministic populations.

MF formulations for heterogeneous networks subject to extrinsic noise have been examined in the context of the *circular cumulants* expansion [43, 45, 53, 54]. However, any finite truncation of this expansion leads to a divergence of the population firing rate [54]. Our formulation does not suffer of these strong limitations and even the definitions of the macroscopic observables are not modified by considering higher order corrective terms [47].

In order to clarify the limits of applicability of our formulation, we have introduced in [47] a reference noise scale up to which a good quantitative agreement between network simulations and the MF model (10) should be expected. Furthermore, preliminary results suggest that our approach can be extended also to noisy homogenous networks displaying sustained firing activities [47].

Potentially, the introduced framework can be fruitfully applied to generalize previous results obtained for many body systems with LD heterogeneities [2, 4, 55].

## Acknowledgements

We acknowledge stimulating discussions with Lyudmila Klimenko, Gianluigi Mongillo, Arkady Pikovsky, and Antonio Politi. The development of the basic theory of pseudo-cumulants was supported by the Russian Science Foundation (Grant No. 19-42-04120). A.T. and received financial support by the Excellence Initiative I-Site Paris Seine (Grant No. ANR-16-IDEX-008), by the Labex MME-DII (Grant No. ANR-11-LBX-0023-01), and by the ANR Project ERMUNDY (Grant No. ANR-18-CE37-0014), all part of the French program Investissements d’Avenir.

## Supplemental Material

### CHARACTERISTIC FUNCTION AND PSEUDO-CUMULANTS

Here we report in full details the derivation of the model (10), already outlined in the Letter, in terms of the characteristic function and of the associated pseudo-cumulants. In particular, the characteristic function for *V*_***x***_ is defined as

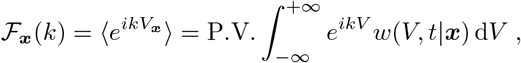

which for a Lorentzian distribution becomes:

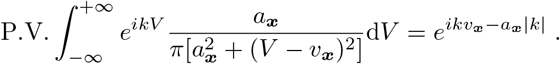

In order to derive the FPE in the Fourier space, let us proceed with a more rigourous definition of the characteristic function, namely

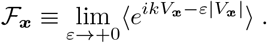

Therefore by virtue of the FPE (Eq. (3) in the Letter) the time derivative of the characteristic function takes the form

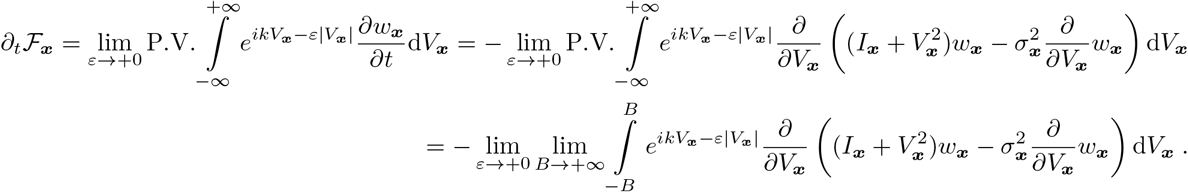

Performing a partial integration, we obtain

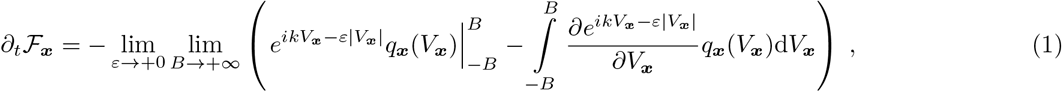

where the probability flux for the ***x***-subpopulation is defined as

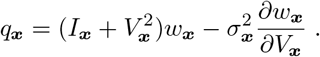

As the membrane potential, once it reaches the threshold +*B*, is reset to −*B* this sets a boundary condition on the flux, namely *q*_***x***_(*B*) = *q*_***x***_(−*B*) for *B* → +∞; therefore,

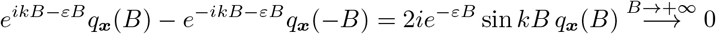

and the first term in Eq. (1) will vanish, thus the time derivative of the characteristic function is simply given by

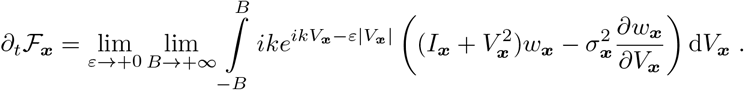

Hence, after performing one more partial integration for the remaining *V*_****x****_-derivative term, we obtain

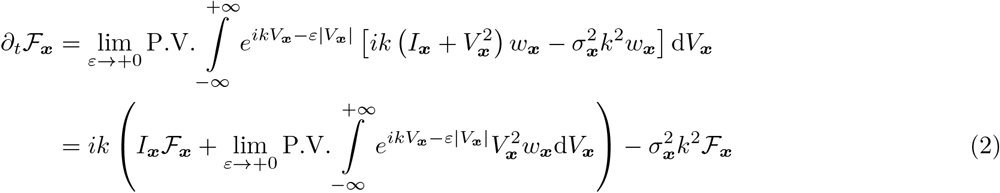

and finally

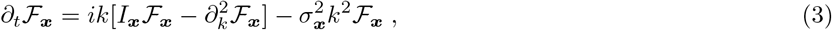

which is Eq. (4) in the Letter.

Under the assumption that 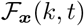 is an analytic function of the parameters ***x*** one can calculate the average characteristic function for the population 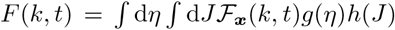 and the corresponding FPE via the residue theorem, with the caution that different contours have to be chosen for positive (upper half-planes of complex *η* and *J*) and negative *k* (lower half-planes). Hence, the FPE is given by

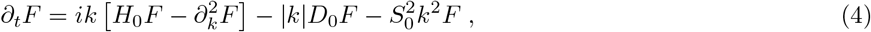

where *H*_0_ = *I*_0_ + *η*_0_ + *J*_0_*r*, *D*_0_ = Δ_*η*_ + Δ_*J*_*r* and 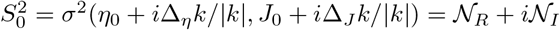.

For the logarithm of the characteristic function, *F(k)* = *e*^Φ(*k*)^, one obtains the following evolution equation

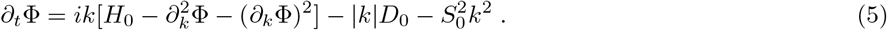

In this context the Lorentzian Ansatz amounts to set Φ_*L*_ = *ikv* − *a*|*k*| [1], by substituting Φ_*L*_ in (5) for *S*_0_ = 0 one gets

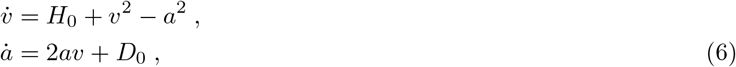

which coincides with the two dimensional mean-field model found in [2] with *r* = *a*/*π*.

In order to consider deviations from the Lorentzian distribution, we analyse the following general polynomial form for Φ:

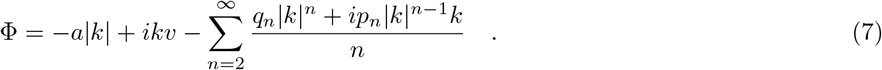

The terms entering in the above expression are dictated by the symmetry of the characteristic function 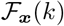 for real-valued *V*_***x***_, which is invariant for a change of sign of *k* joined to the complex conjugation. For this characteristic function neither moments, nor cumulats can be determined [3].

Hence, we can choose the notation in the form which would be most optimal for our consideration. Specifically, we introduce Ψ = *k*∂_*k*_Φ,

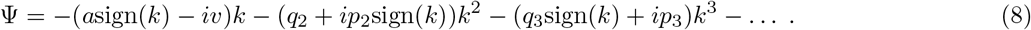

Please notice that

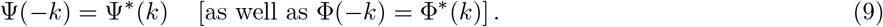

In this context Eq. (5) becomes

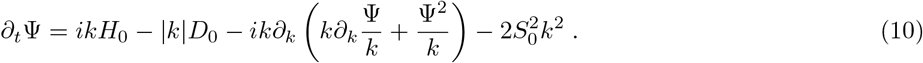

It is now convenient to introduce the *pseudo-cumulants*, defined as follows:

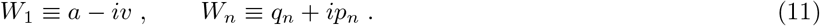

From Eq. (10) we can thus obtain the evolution equation for the pseudo-cumulants *W_m_*, namely

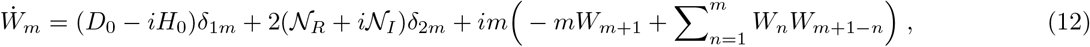

where for simplicity we have assumed *k* > 0 and employed the property (9). Moreover, we have omitted the *kδ(k)* contribution, since it vanishes.

The evolution of the first two pseudo-cumulant reads as:

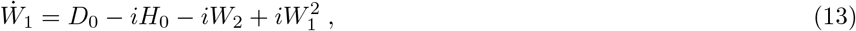

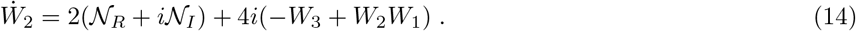

Or equivalently

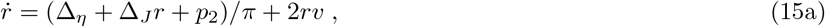

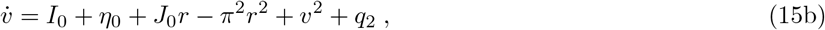

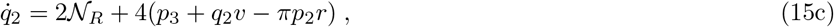

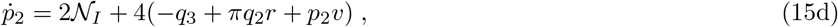

which is Eq. (10) in the Letter.

### FIRING RATE AND MEAN MEMBRANE POTENTIAL FOR PERTURBED LORENTZIAN DISTRIBUTIONS

In the following we will demonstrate that the definitions of the firing rate *r* and of the mean membrane potential *v* in terms of the PDF *w(V, t)*, namely:

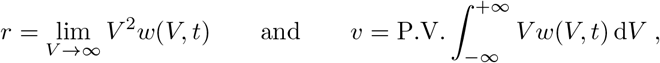

obtained in [2] for a Lorentzian distribution, are not modified even by including in the PDF the correction terms {*q_n_*, *p_n_*}.

The probability density for the membrane potentials *w(V, t)* is related to the characteristic function *F(k)* via the follwoing anti-Fourier transform

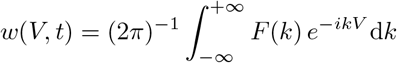

with *F(k)* = *e*^Φ(*k*)^. By considering the deviations of Φ(*k*) from the Lorentzian distribution up to the second order in *k*, we have

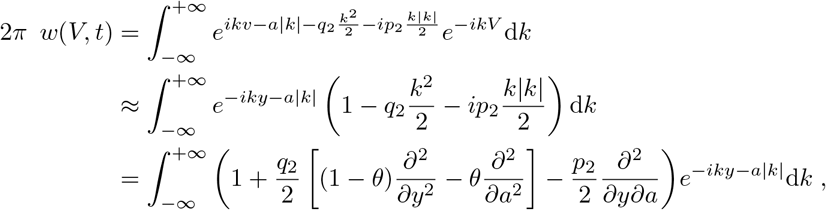

where *y* = *V* − *v* and *θ* is an arbitrary parameter. Thus one can rewrite

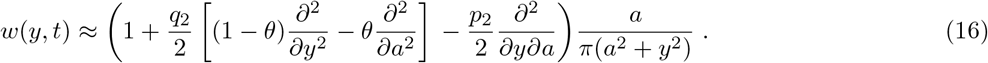

From the expression above, it is evident that *q*_2_ and *p*_2_, as well as the higher-order corrections, do not modify the firing rate definition reported in [2] for the Lorentzian distribution, indeed

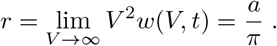

Let us now estimate the mean membrane potential by employing the PDF (16), where we set the arbitrary parameter
*θ* to zero without loss of generality, namely

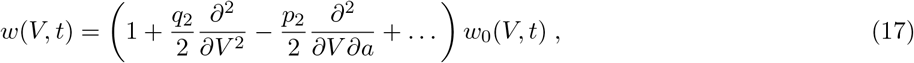

where *w*_0_(*V, t*) = *π*^−1^*a*/[*a*^2^ + (*V* − *v*)^2^]. The mean membrane potential is given by

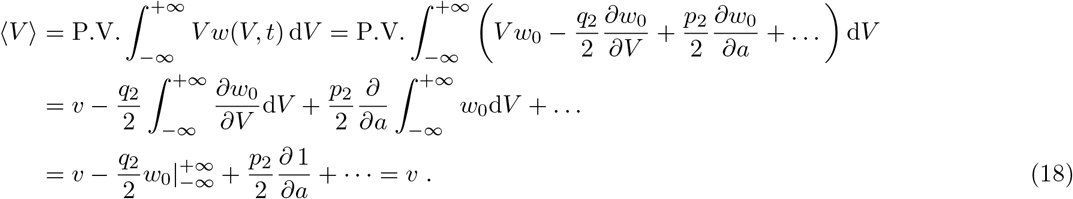

All the higher-order corrections entering in *w(V, t)*, denoted by (…) in (18), have the form of higher-order derivatives of *w*_0_ with respect to *V* and a; therefore they yield a zero contribution to the estimation of 〈*V*〉. Thus, Eq. (18) is correct not only to the 2nd order, but also for higher orders of accuracy. We can see that the interpretation of the macroscopic variables *a* = *πr* and *v* = 〈*V*〉 in terms of the firing rate and of the mean membrane potential entering in Eq. (12) or Eqs. (15a)–(15d) remains exact even away from the Lorentzian distribution.

### SMALLNESS HIERARCHY OF THE PSEUDO-CUMULANTS

Eq. (12) for *m*> 1 can be recast in the following form

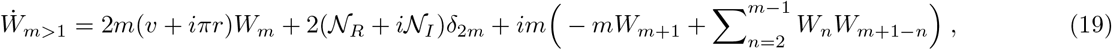

where *W_m_* is present only in the first term of the right-hand side of the latter equation.

Let us now understand the average evolution of *W_m_*, *m* > 1. In particular, by dividing Eq. (15a) by *r* and averaging over time, one finds that

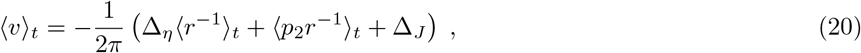

where 〈…〉_*t*_ denotes the average over time and where we have employed the fact that the time-average of the time-derivative of a bounded process is zero, i.e. 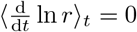. Since *r(t)* can be only positive, 〈*v*〉_*t*_ will be strictly negative for a heterogeneous population (with Δ_*η*_ ≠ 0 and/or Δ_*J*_ ≠ 0) in the case of nonlarge deviations from the Lorentzian distribution, i.e., when *p*_2_ is sufficiently small. In particular, for asynchronous states *v* = 〈*v*〉_*t*_, hence, Eq. (20) yields a relaxation dynamics for *W_m_* under forcing by *W_m+1_* and *W_1_*,…, *W_m−1_*; by continuity, this dissipative dynamics holds also for oscillatory regimes which are not far from the stationary states.

Let us explicitly consider the dynamics of the equations (19) for *m* = 2, 3, 4, namely:

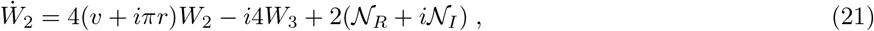

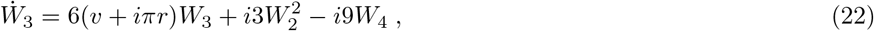

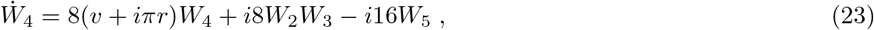

Let us first see how the attractivity of the Lorentzian distribution in the absence of noise follows from these equations. For 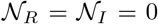, we consider a small deviation from the Lorentzian distribution such that |*W_n_*| < *Cε*^*n*−1^, where *C* is some positive constant and *ε* ≪ 1 is a smallness parameter. In this case, from Eq. (21) one observes that *W*_2_ tends to ~ *W*_3_, while from Eq. (22), 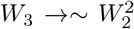. Here *W*_4_ is neglected as for the initial conditions one finds 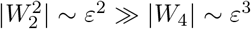 and below we will see that similar relation remains valid over time. Therefore 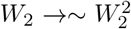, which means that *W*_2_(*t* → +∞) → 0. Further, from Eq. (23), 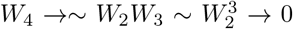. Here *W*_5_ is neglected as for the initial conditions one finds |*W*_2_*W*_3_| ~ *ε*^3^ ≫ |*W*_5_| ~ *ε*^4^. In the course of evolution *W*_4_ tends to 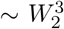, while *W*_3_ tends to 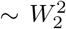, one can similarly show that 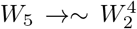, and so forth. Hierarchy 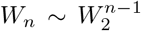 is similar to *W_n_* ~ *ε*^*n*−1^ and allows us to neglect *W*_4_ in 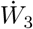, *W*_5_ in 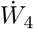, and so forth, not only for the initial stage of the dynamics, but also at a later time. Thus, in the absence of noise, the system tends to a state *W*_1_ ≠ 0, *W_m>1_* = 0 (at least from a small but finite vicinity of this state). This tells us that the Lorentzian distribution is an attractive solution in this case.

In the presence of noise, by assuming that 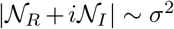, a similar analysis of Eqs. (21)–(23) yields |*W*_2_| →~*σ*^2^, 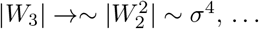,

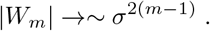

The above scaling is well confirmed by the data reported in Fig. 1. Therefore, a two-element truncation (13)–(14) of the infinite equation chain (12) is well justified as a first significant correction to the Lorentzian distribution dynamics. Presumably, this might also hold for some regimes in homogeneous populations (where Δ_*η*_ = Δ_*J*_ = 0), even thought the heuristic explanation we provide here heavily relays on the positivity of (Δ_*η*_〈*r*^−1^〉_*t*_ + Δ_*J*_) in Eq. (20).

**FIG. 1.**
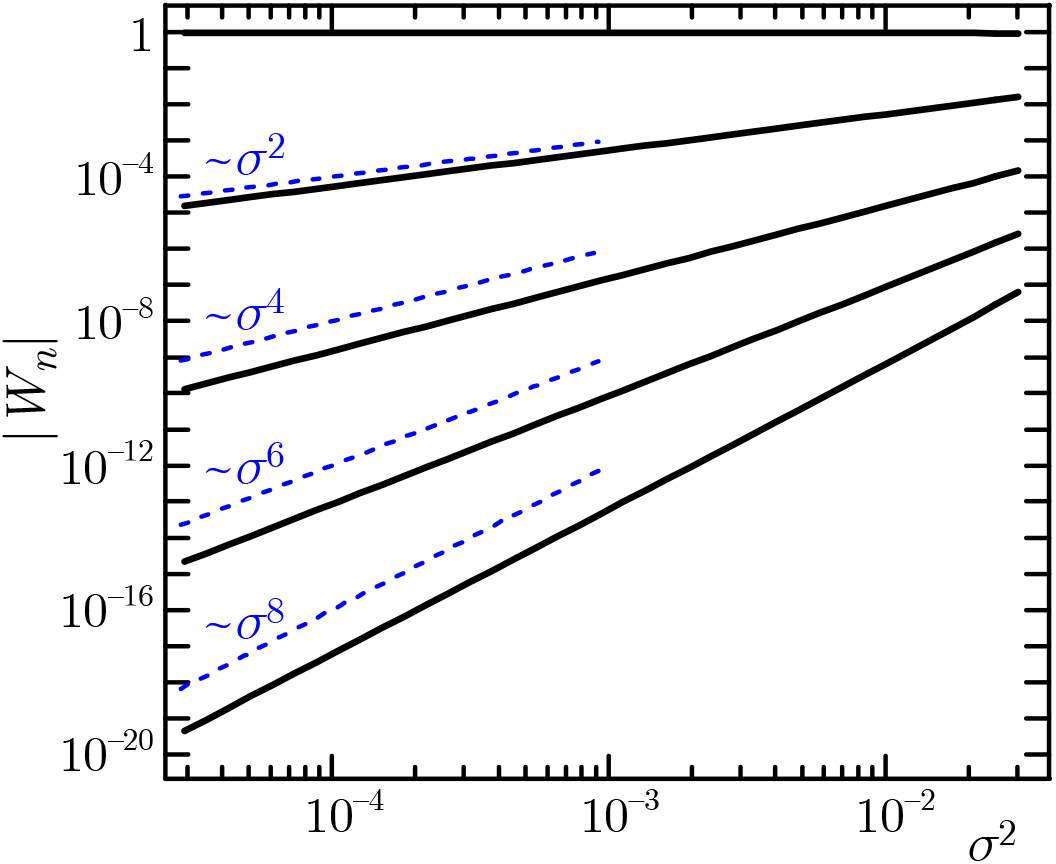
Modulus of the pseudo-cumulants |*W_n_*| versus the noise variance *σ*^2^ for *n* ranging from 1 to 5 from top to bottom. The pseudo-cumulants are estimated by integrating Eq. (10) in the Letter with extended precision (30 digits) and by limiting the sum to the first 100 elements. Other parameters: *I*_0_ = 0.1, *η*_0_ = −1, *J*_0_ = 1, Δ*η* = 0.1, and Δ_*J*_ = 0.1.

### CONVENTIONAL AND PSEUDO- CUMULANTS

#### The relationships between conventional and pseudo- cumulants

Let us discuss the relationships existing between conventional and pseudo-cumulant representations. The characteristic function *F(k)* and its logarithm Φ(*k*) of any *real-valued* random variable *V* must obey the symmetry properties *F(k)* = *F*\*(−*k*) and Φ(*k*) = Φ*(−*k*). Hence, the expression (7) for the function Φ(*k*) in terms of pseudo-cumulants is the most general one respecting such symmetry

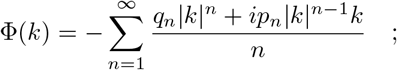

where we set *q*_1_ = a and *p*_1_ = −*v*.

Thus, the pseudo-cumulants can be expressed as derivative of Φ(*k*), as follows:

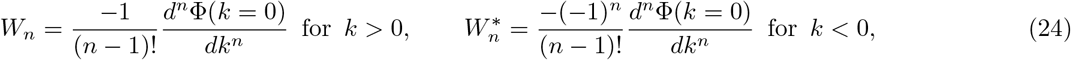

where 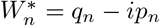 is the complex conjugate of *W_n_*.

However, the expression (7) does not guarantee the existence of conventional cumulants and moments. The existence of the *n*-th moment *M_n_* = 〈*V^n^*〉 requires that the PDF *w(V)* will decay faster than 1/|*V*|^*n*+1^ for *V* → ±∞. If the *n*-th moment is finite, the derivative 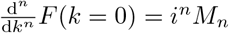 exists and is continuous (as well as all lower-order derivatives). The finiteness of 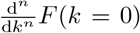 forbids the existence of all the terms |*k*|^2*m*−1^ with odd (2*m*−1) ≤ *n* and |*k*|^2*m*−1^*k* with even 2*m* ≤ *n*, meaning that only smooth terms *k*^2*m*−1^ and *k*^2*m*^ are present up to the *n*-th power (the *k^n^*-term included). Therefore, in this case we will have

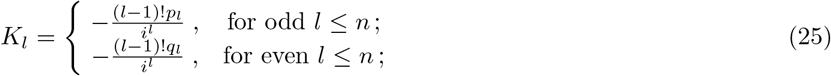

where *K_l_* is the *l*-th order *conventional cumulant*, defined from the expansion 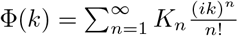.

To summarize, if the distribution *w(V)* decays as 1/|*V*|^*n*+1+α^ with 0 < α ≤ 1 then we have the following situation. The expansion of Φ(*k*) contains for the orders up to *l* ≤ *n* only the conventional-cumulant part, *K*_1_ +…+*K_l_(ik)^l^/l*!+…, and the pseudo-cumulants are purely real (imaginary) for even (odd) *l* in agreement with (25). For the orders *l* > *n*, conventional moments and cumulants diverge and the pseudo-cumulants *W_l_* have generally both the real *q_l_* and imaginary *p_l_* parts given by (24).

In particular, if the distribution *w(V)* decays faster that any power law (e.g., exponentially fast) then all the moments are finite and all pseudo-cumulants have only the conventional-cumulant part.

#### Geometric interpretation of the second pseudo-cumulant

The small-*k* behavior of the characteristic function *F(k)*, which is the Fourier transform of the PDF *w(V)*, represents the large-scale properties of *w(V)*. In particular, the leading part of the expansion reads as

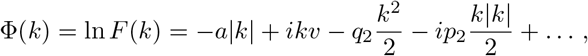

where the terms ~ *k* characterize the ‘Lorentzian’ profile of the distribution; while the second order terms ~ *k*^2^ represent the leading corrections to the distribution for large *V*.

In particular, *q*_2_ = Re(*W*_2_) can be interpreted as an analogue of the *kurtosis* for distorted Gaussian distributions. Indeed, Eq. (17) reads as 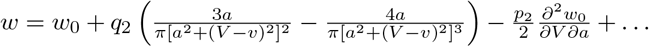 and *q*_2_ > 0 implies an increase. of the deviations from *w*_0_ at large |*V*|.

The *p*_2_-term is odd in *k* and therefore represents the asymmetry in *w(V)* between *V* and *−V* ; this is derived from the fact that *F(-k)* = 〈*e*^*ik(-V)*^〉. Moreover, from Eq. (16) one can see, that the median of the distribution *w(V)* is not affected by the *p*_2_-term. This can be easily shown by evoking the expression for *w(V)* reported in (17) and by noticing that, according to the definition of a median, the integral of the Lorentzian distribution *w*_0_(*V*) over the half-axis *V* ∈[*v*; +∞) equals 1/2. Indeed, the following integral vanishes

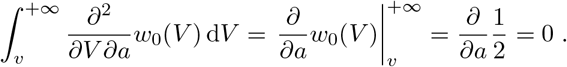

Hence, the integral of *w(V)* over the half-axis *V* ∈[*v*; +∞) is still 1/2 and not modified by the *p*_2_-term. Similarly the median is not affected by the *q*_2_-term, as 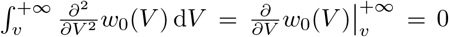. Therefore, *v* remains the median of the distribution. Thus, *p*_2_ = Im(*W*_2_) measures the asymmetry of *w(V)* for a fixed median value given by *v* = −Im(*W*_1_) and therefore it can be interpreted as the *skewness* of the distorted distribution.

In the specific case of QIFs, the integrals over *V* are defined as the principal value ones and the mean value of *V* can be calculated also for a distribution with Lorentzian tails. Thus, here one can speak not only of the median—this interpretation will be valid universally—but also of the mean value; as one can see from Eq. (18), the p_2_-term does not shift the population-mean value 〈*V*〉.

### A REFERENCE SCALE FOR THE NOISE

Let us reconsider the MF equations (15), in particular equation (15a) divided by *r* and averaged over time yields

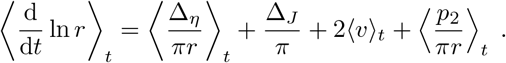

As we have already mentioned 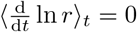 and *r* is always positive, therefore for *p*_2_ ≪ 1, one finds 〈*v*〉_*t*_ < 0. To clarify the relevance of the noise term, one can rewrite Eqs. (15c)–(15d) as

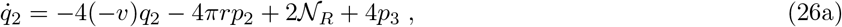

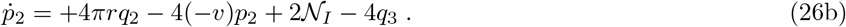

The latter system yields on average a linearly decaying dynamics with average decay rate −4〈*v*〉_*t*_ plus a counterclock-wise rotation on the plane (*q*_2_, *p*_2_) with instantaneous angular velocity 4π*r* and with constant driving terms given by 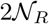 and 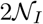 plus small (*q*_3_, *p*_3_) corrections. Therefore, the effect of noise is fundamental in order to obtain non-vanishing values for the terms *q*_2_ and *p*_2_.

Indeed, this becomes evident in the stationary case, where by neglecting (*q*_3_, *p*_3_) in Eqs. (26), one obtains

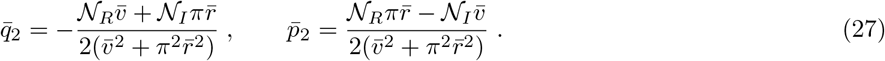

In the case of globally coupled network with additive noise of amplitude *σ*, where 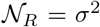 and 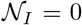, the terms 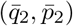 are directly proportional to the noise variance *σ*^2^, as already shown in Fig. 1 (c-d) in the Letter.

An estimation of a reference value for the noise strength can be obtained by considering the effect of the stationary terms (27) on the evolution of the firing rate and of the mean membrane potential given by Eqs. (15a)–(15b). In this case we can give a clear physical interpretation of the stationary corrections 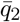 and 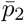. They can be interpreted as a measure of an additional source of heterogeneity in the system induced by the noise. To be more precise, 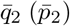 can be considered as a modification of the mean input current in (15b) (of the width of the distribution of the heterogeneities in (15c)) and therefore it should be compared with the median of the effective input current *I*_0_ + *η*_0_ + *J*_0_*r* (with the HWHM of the effective input currents Δ_*η*_ + Δ_*J*_*r*) appearing in the same equation.

A first reference value for the noise strenght 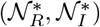 can be estimated from (15a) by setting 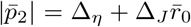, thus obtaining

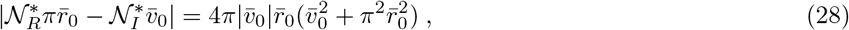

where 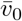 and 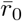 are the time-independent solution of Eqs. (15a)–(15b) for 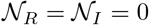.

From (15b) a second reference scale for the noise 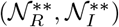 can be derived by setting 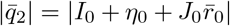, thus obtaining

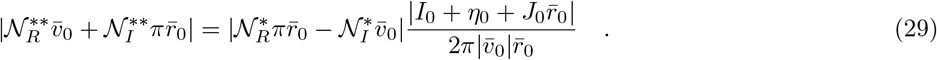

The solution of the above equation, which would define the second scale, is proportional to the solution of (28) which will set 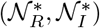, therefore it is justified to consider only the latter ones as the reference values for the noise amplitude. In the following we will estimate this reference noise for the specific cases considered in the Letter.

#### Globally coupled network with extrinsic noise

For the globally coupled network with addtive noise we have 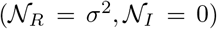, therefore the reference noise strenght *σ*_*_ is given by

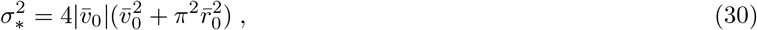

and the second noise scale *σ*_**_ is obtained from (29)

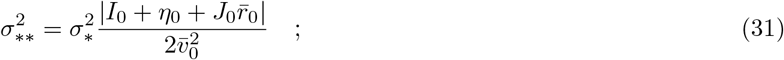

which confirms that we can limit to consider as a scale for the noise *σ*_*_, since the second noise variance value is proportional to the first one.

Let us restrict our analysis to Δ_*η*_ = 0, where 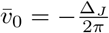, and 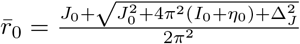. In this case we have an explicit expression for the reference noise scale, i.e.

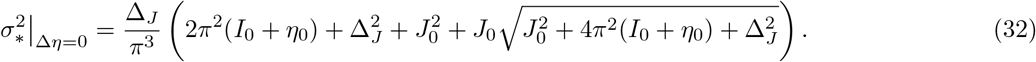

In particular, for the parameters employed in Fig. 1 (a-e) in the Letter, i.e. *I*_0_ = 0.0001, *η*_0_ = 0, *J*_0_ = −0.1, Δ_*J*_ = 0.1, one obtains *σ*_*_ ≈ 0.00458, while for those employed in Fig. 1 (f-g) in the Letter, i.e. *I*_0_ = 0.38, *η*_0_ = 0 *J*_0_ = −6.3, Δ_*J*_ = 0.01, one finds *σ*_*_ ≈ 0.014. In Fig. 1 of the Letter we have employed the above values *σ*_*_ to rescale the noise amplitudes as 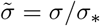. The correctness of the choice of this reference values is confirmed from the fact that in the asynchronous state, analysed in Fig. 1 (a-b), the deviations from the MPR results (magenta dashed lines) become evident for 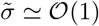.

#### Sparse networks exhibiting endogenous fluctuations

For the sparse deterministic network under the Poissonian approximation for the input spike trains, we can write 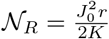 and 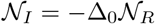, with Δ_*J*_ = Δ_0_|*J*_0_|. In this case, the reference scale for the noise is simply given by

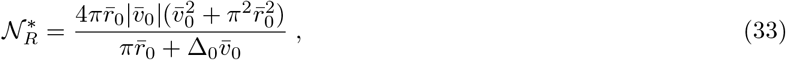

since 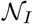 is directly proportional to 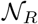.

Once more we consider the case Δ_*η*_ = 0, where we have an explicit expression for the mean membrane potential 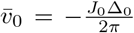 and for the firing rate 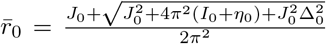. More specifically, for the parameters employed in Fig. 2 in the Letter, namely *K* = 5000, Δ_0_ = 0.01 and *η* = 0, we obtain for a large coupling value *J*_0_ = −5 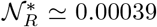 for *I*_0_ = 0.19 as in panel (b) and 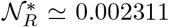 for *I*_0_ = 0.50 as in panel (c). In these two specific cases, we measured the corresponding average firing rates and from these values we have obtained an estimate of the average 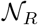 and of the corresponding rescaled noise amplitude 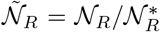. In particular, for *J*_0_ = −5 we found 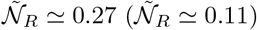 for the case reported in panel (b) (panel (c)). This difference in the 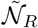-values explains why for corresponding synaptic coupling the quantitative agreement between network simulations and MF results is worst in panel (b).

### NOISY HOMOGENEOUS POPULATIONS

The Ott-Antonsen approach [4], as well as the MPR mean-field model [2], have been derived for heterogeneous deterministic systems and it is known that the corresponding reduced manifolds are no more attractive for homogeneous populations [5].

Our approach has been developed for heterogenous noisy populations: an important question is if and when it can be extended to the case of populations of identical element. This specific point goes beyond the scopes of this Letter and it will be addressed in a future publication [6]. However, here report preliminary analyses showing that there are situations where our MF model reproduces perfectly the network dynamics even in completely homogenous situations. Two examples are shown in Fig. 2 (a-d) for a fully coupled network with additive Gaussian noise for two different noise amplitudes. In these specific examples the external DC current is initially set to *I*_0_ = 2, where the system reveals an asynchronous dynamics, corresponding to a stable focus in the MF. At a time *t* = 20 current *I*_0_ is increased to a value 4, where the system displays COs, and maintained at such value for a certain time interval and then restored to the initial value. As evident from the figures, the MF evolution is in perfect agreement with the network dynamics, apart finite size fluctuations, and it is even able to capture the relaxation oscillations towards the stable focus at times *t*> 50.

In Fig. 2 (e) we consider a cut at constant noise amplitude *σ* = 0.006 in the phase diagram reported in Fig. 1 (e) of the Letter. In particular, we examine the evolution of the network and MF dynamics by decreasing adiabatically Δ*J* from an initial finite value (Δ*J* = 0.1) to a vanishingly small value of Δ*J*, then we increase again adiabatically the parameter back to the initial value. For the network we can reach Δ*J* = 0 (the homogeneous case), while the MF exhibits diverging solutions for Δ*J* → 0. However, the MF captures the Hopf sub-critical bifurcation from the asynchronous dynamics to COs at Δ*J*_*HB*_ ≃ 0.0089 (black solid line in Fig. 1 (e) of the Letter) as well as the saddle-node of limit cycles at Δ*J_SN_* ≃ 0.06255 (red solid line in Fig. 1 (e) of the Letter) displayed also by the network, apart finite size corrections (as shown in Fig. 2 (e)). Furthermore, the standard deviation of the mean membrane potential ∑_*v*_ are reasonably well reproduced down to Δ*J* ≃ 0.02, i.e. for systems that we can consider *de facto* as homogenous due to the quite large value of the median of the synaptic coupling, namely *J*_0_ = −6.3. Therefore, it is true that in this case the MF gives diverging solutions in the homogenous case, however the homogenous solutions are essentially indistinguishable from the heterogeneous one at Δ*J* = 0.02, where the MF still gives reasonable results.

From our preliminary analyses [6], it emerges that the homogeneous case is better captured by a MF approach for not vanishingly small values of the firing rate. This seems consistent with our findings in the present case. Indeed, in Fig. 2 (c) and (d) the population firing rate is *r* ≃ 0.5 − 1.0, while for the case reported in Fig. 2 (e) *r* ≃ 0.05 − 0.08, i.e. much smaller. A *theoretical* criterion for the applicability of the MF formulation can be formulate as follows: if the population-mean firing rate (*or* other mean fields driving the macroscopic dynamics) as a function of a small parameter (e.g., the noise intensity) can be represented by a power series, the approach can be safely employed in for noisy homogenous populations. As an example, if the firing rate follows a law like *σ*^*n*^ exp(−*A*/*σ*^2^) such a dependence cannot be represented by its power series, which formally reads as *σ*^*n*^(0 + 0.*σ*^2^ +0.*σ*^4^ +…) [7]. Formal application of the pseudo-cumulant approach to this case will yield *r* = 0.

**FIG. 2.**
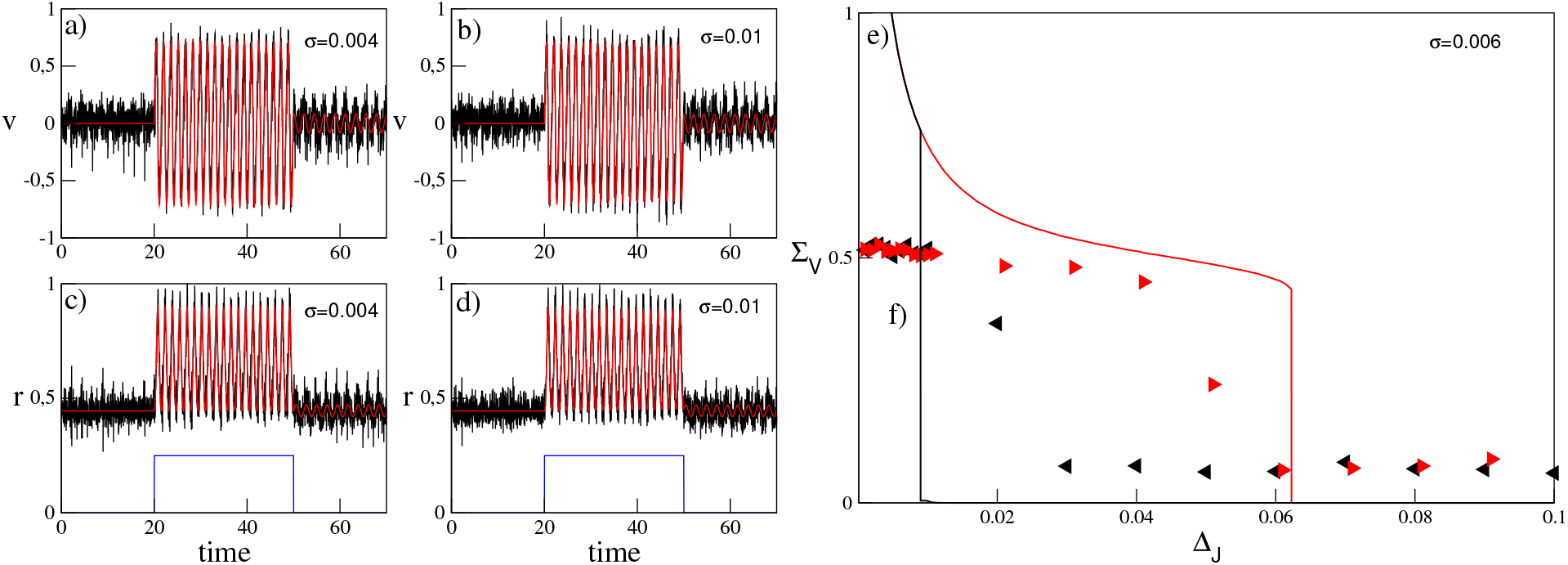
Homogeneous Globally Coupled Network subject to Additive Noise. (a-d) Red (black) solid line show the time course of *v* (a-c) and r (b-d) of the MF (of the network with *N* = 10000). Panels (a) and (c) refer to *σ* = 0.004, panels (b) and (d) to *σ* = 0.01. The external current is *I*_0_ = 4 for time ∈ [20, 50] and *I_0_* = 2 otherwise, *I*_0_ is shown as blue solid line in panels (c) and (d) after a suitable rescaling. Other parameters: *J*_0_ = −0.1 and *η*_0_ = Δ*J* = Δ_*η*_ = 0. (e) Standard deviation ∑_*v*_ versus Δ_*J*_ a noise amplitudes *σ* = 0.00Δ Lines (symbols) refer to MF (network) results: solid red (black) lines and right (left) triangles are obtained by increasing (decreasing) Δ*J*. Other parameters are as in Fig.1 (f) of the Letter: namely, *I*_0_ = 0.38, *J*_0_ = −6.3 and *η*_0_ = Δ*η* = 0.

## References

[1] V. M. Zolotarev, One-dimensional stable distributions, Translations of Mathematical Monographs, vol. 65 (1986).

[2] E. Yakubovich, Soviet Physics JETP 28, 160.

[3] W. E. Lamb Jr, Physical Review 134, A1429 (1964).

[4] P. Lloyd, Journal of Physics C: Solid State Physics 2, 1717 (1969).

[5] P. W. Anderson, Physical review 109, 1492 (1958).

[6] M. Rabinovich and D. Trubetskov, “Oscillations and waves: In linear and nonlinear systems (vol. 50),” (1989).

[7] J. D. Crawford, Journal of Statistical Physics 74, 1047 (1994).

[8] D. S. Goldobin and A. Pikovsky, Physical Review E 71, 045201 (2005).

[9] D. S. Goldobin and A. V. Dolmatova, Communications in Nonlinear Science and Numerical Simulation 75, 94 (2019).

[10] E. Ott and T. M. Antonsen, Chaos: An Interdisciplinary Journal of Nonlinear Science 18, 037113 (2008).

[11] E. Ott and T. M. Antonsen, Chaos: An interdisciplinary journal of nonlinear science 19, 023117 (2009).

[12] A. T. Winfree, Journal of theoretical biology 16, 15 (1967).

[13] Y. Kuramoto, Chemical oscillations, waves, and turbulence (Courier Corporation, 2003).

[14] S. Watanabe and S. H. Strogatz, Physica D: Nonlinear Phenomena 74, 197 (1994).

[15] S. A. Marvel and S. H. Strogatz, Chaos: An Interdisciplinary Journal of Nonlinear Science 19, 013132 (2009).

[16] A. V. Dolmatova, D. S. Goldobin, and A. Pikovsky, Physical Review E 96, 062204 (2017).

[17] V. Klinshov and I. Franović, Physical Review E 100, 062211 (2019).

[18] I. V. Tyulkina, D. S. Goldobin, L. S. Klimenko, I. S. Poperechny, and Y. L. Raikher, Philosophical Transactions of the Royal Society A 378, 20190259 (2020).

[19] T. B. Luke, E. Barreto, and P. So, Neural Computation 25, 3207 (2013).

[20] C. R. Laing, Physical Review E 90, 010901 (2014).

[21] E. Montbrió, D. Pazó, and A. Roxin, Phys. Rev. X 5, 021028 (2015).

[22] F. Devalle, A. Roxin, and E. Montbrió, PLoS computational biology 13, e1005881 (2017).

[23] A. Byrne, M. J. Brookes, and S. Coombes, Journal of computational neuroscience 43, 143 (2017).

[24] G. Dumont, G. B. Ermentrout, and B. Gutkin, Physical Review E 96, 042311 (2017).

[25] F. Devalle, E. Montbrió, and D. Pazó, Physical Review E 98, 042214 (2018).

[26] H. Schmidt, D. Avitabile, E. Montbrió, and A. Roxin, PLoS computational biology 14, e1006430 (2018).

[27] S. Coombes and A. Byrne, in Nonlinear Dynamics in Computational Neuroscience, edited by F. Corinto and A. Torcini (Springer, 2019) pp. 1–16.

[28] B. Pietras, F. Devalle, A. Roxin, A. Daffertshofer, and E. Montbrió, Physical Review E 100, 042412 (2019).

[29] G. Dumont and B. Gutkin, PLoS Computational Biology 15, e1007019 (2019).

[30] A. Ceni, S. Olmi, A. Torcini, and D. Angulo-Garcia, Chaos: An Interdisciplinary Journal of Nonlinear Science 30, 053121 (2020).

[31] M. Segneri, H. Bi, S. Olmi, and A. Torcini, Front. Comput. Neurosci. 14 (2020).

[32] H. Taher, A. Torcini, and S. Olmi, PLOS Computational Biology 16, 1 (2020).

[33] E. Montbrió and D. Pazó, Physical Review Letters 125, 248101 (2020).

[34] E. R. Kandel, J. H. Schwartz, T. M. Jessell, S. Siegelbaum, A. J. Hudspeth, and S. Mack, Principles of neural science, Vol. 4 (McGraw-hill New York, 2000).

[35] C. Van Vreeswijk and H. Sompolinsky, Science 274, 1724 (1996).

[36] J. Barral and A. D. Reyes, Nature neuroscience 19, 1690 (2016).

[37] C. Geisler, N. Brunel, and X.-J. Wang, Journal of neurophysiology 94, 4344 (2005).

[38] N. Brunel and V. Hakim, Neural computation 11, 1621 (1999).

[39] N. Brunel, Journal of computational neuroscience 8, 183 (2000).

[40] M. Mattia and P. Del Giudice, Physical Review E 66, 051917 (2002).

[41] E. S. Schaffer, S. Ostojic, and L. F. Abbott, PLoS Comput Biol 9, e1003301 (2013).

[42] T. Schwalger, M. Deger, and W. Gerstner, PLoS computational biology 13, e1005507 (2017).

[43] B. Pietras, N. Gallice, and T. Schwalger, Phys. Rev. E 102, 022407 (2020).

[44] M. di Volo and A. Torcini, Physical review letters 121, 128301 (2018).

[45] I. Ratas and K. Pyragas, Physical Review E 100, 052211 (2019).

[46] G. B. Ermentrout and N. Kopell, SIAM Journal on Applied Mathematics 46, 233 (1986).

[47] See Supplemental Material for a detailed derivation of the MF model (10) and of the expressions of r and v for perturbed LDs, for an estimation of the scaling of the |*W_n_*| with the noise amplitude, for the relations between conventional and pseudo-cumulants, for the derivation of a reference noise scale, and for a preliminar analysis of the homogenous noisy case.

[48] The Fourier transform of the Lorentzian distribution is P. V. 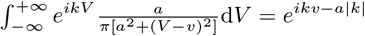

[49] E. Lukacs, Characteristic functions (Griffin, 1970).

[50] H. Bi, M. Segneri, M. di Volo, and A. Torcini, Physical Review Research 2, 013042 (2020).

[51] A. Renart, R. Moreno-Bote, X.-J. Wang, and N. Parga, Neural computation 19, 1 (2007).

[52] H. R. Wilson and J. D. Cowan, Biophysical journal 12, 1 (1972).

[53] I. V. Tyulkina, D. S. Goldobin, L. S. Klimenko, and A. Pikovsky, Physical review letters 120, 264101 (2018).

[54] D. S. Goldobin and A. V. Dolmatova, Physical Review Research 1, 033139 (2019).

[55] I. Lifshitz, S. Gredeskul, and L. Pastur, New York (1988).

## References

[1] The Fourier transform of the Lorentzian distribution is P.V. 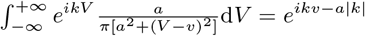

[2] E. Montbrió, D. Pazó, and A. Roxin, Phys. Rev. X 5, 021028 (2015).

[3] E. Lukacs, Characteristic functions (Griffin, 1970).

[4] E. Ott and T. M. Antonsen, Chaos: An Interdisciplinary Journal of Nonlinear Science 18, 037113 (2008).

[5] A. Pikovsky and M. Rosenblum, Physical review letters 101, 264103 (2008).

[6] M. di Volo, M. Segneri, D. Goldobin, A. Politi, and A. Torcini, in preparation (2021).

[7] Function *σ*^*n*^ exp(−*A*/*σ*^2^) possesses an essential singularity at *σ* = 0 even though it is infinitely smooth at this point.

